# AlphaFind v2: Similarity Search in AlphaFold DB and TED Domains across Structural Contexts

**DOI:** 10.64898/2026.03.10.710735

**Authors:** Terézia Slanináková, Adrián Rošinec, Jakub Čillík, Aleš Křenek, Katarina Gresova, Jana Porubská, Eva Maršálková, Jaroslav Olha, David Procházka, Lukáš Hejtmánek, Vlastislav Dohnal, Karel Berka, Radka Svobodová, Matej Antol

**Author notes:** Corresponding author. Matej Antol.

## Abstract

The availability of large-scale protein structure collections enables structure-based analysis of their function and evolution beyond what is possible from sequence alone. However, applying three-dimensional structure comparison at scale remains computationally demanding and limits practical exploration of large experimental and predicted collections. This creates a need for fast, structure-based search methods that retain biological relevance while enabling large-scale exploration. In this paper, we present AlphaFind v2, an application for finding structurally similar proteins in the AlphaFold Database (https://alphafold.ebi.ac.uk/) of predicted structures. AlphaFind v2 uses fast pre-filtering via state-of-the-art protein embeddings that preserve structural information, followed by refinement with US-align. The application presents multiple complementary search modes, including (i) search over full protein chains, (ii) search aware of the AlphaFold pLDDT metric, restricting similarity computation to the most stable and structurally relevant regions, (iii) search over protein domains from the TED database (https://ted.cathdb.info/), and (iv) a multidomain search mode, combining multiple chain-level domain matches within a single score and alignment. The application accepts protein identifiers and returns similar proteins with metrics, rich metadata, and interactive superpositions. AlphaFind v2 additionally allows searching within an organism or CATH label and matches the proteins with experimental structures. AlphaFind v2 is accessible at https://alphafind.ics.muni.cz/.

**Graphical Abstract:** 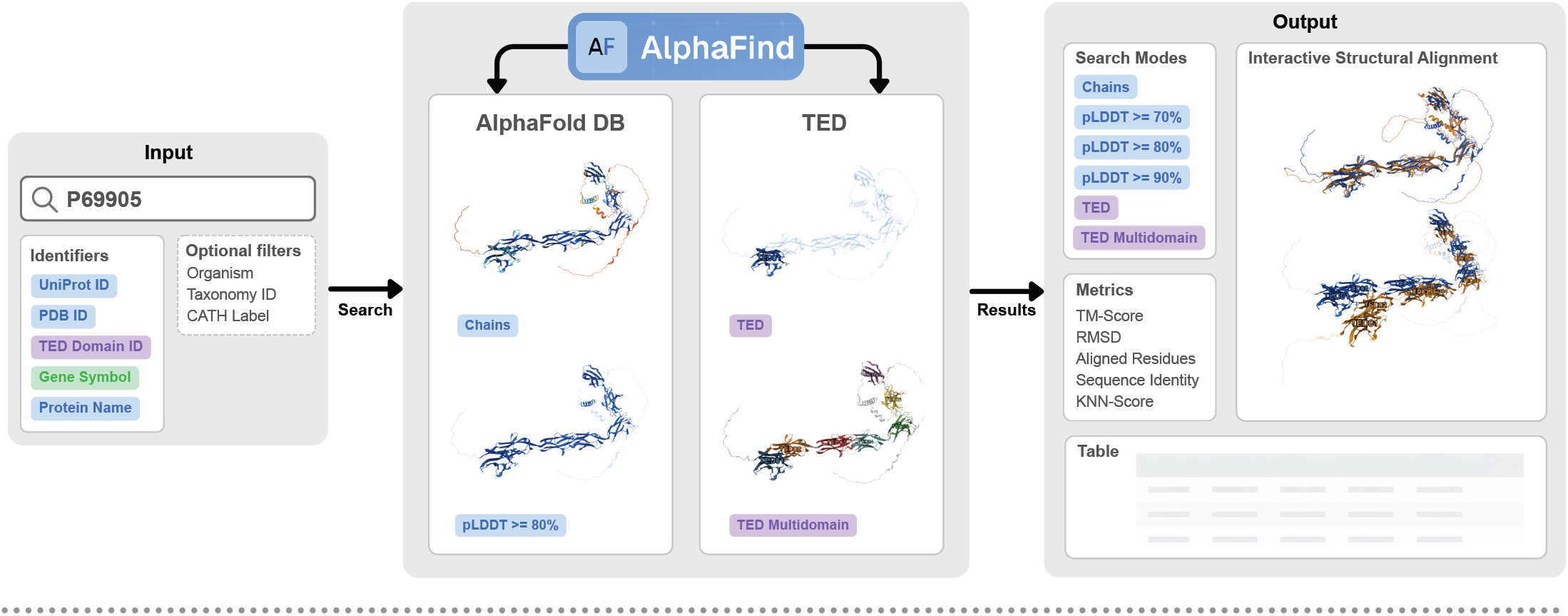

## Introduction

Protein structural data are highly valuable scientific resources that form the foundation of a wide and continuously growing body of research. As of today, the Protein Data Bank, the world’s leading database of experimentally determined protein structures, contains over 227 thousand records (3). Building on these data, the AlphaFold Protein Structure Database (14) now provides more than 240 million predicted protein structures.

Given the scale and diversity of available protein structures, comparing and organizing them in a meaningful way has become a central challenge in structural biology. One of the most prominent approaches is the analysis of protein structural similarity, which provides a powerful lens for understanding biological function and evolutionary relationships. Proteins with similar three-dimensional structures often share functional roles even when their sequences diverge substantially. As the volume of available experimental and predicted structures grew by several orders of magnitude, effective methods for browsing, comparing, and selecting proteins based on structural similarity have become increasingly important.

A variety of approaches exist for quantifying structural similarity, with *alignment-based* measures such as TM-score (16) widely regarded as robust indicators of functional relatedness. However, exact structural alignments are computationally expensive, which limits their applicability in large-scale searches across hundreds of millions of structures.

To address this challenge, recent methods have focused on approximate similarity search strategies that trade exact alignment for speed. Many of these approaches rely on embedding protein structures into lower-dimensional representations that preserve key geometric features (4; 6; 10), enabling fast comparison using standard nearest-neighbor search techniques. Such methods allow large structural databases to be searched efficiently, often serving as a first-stage filter before more precise alignment-based validation. The most prominent examples of search systems using approximate methods include FoldSeek Server (13)(https://search.foldseek.com/) for protein chains and Progres server (4)(https://progres.mrc-lmb.cam.ac.uk/) or Merizo-search (6)(https://bioinf.cs.ucl.ac.uk/psipred) for protein domains. The first version of AlphaFind (9) was introduced in this context as a fast and accessible interface for structure-based search over predicted proteins of the AlphaFold Protein Structure Database.

Here, we introduce AlphaFind v2, which extends the capabilities of its predecessor by enabling direct access not only to full protein chains, but also to their structurally meaningful subsections. In the application, users can restrict searches to regions above a selected confidence threshold of the AlphaFold pLDDT confidence scores, search specific TED domains (7), or use the multidomain search mode. This design allows for flexible, comprehensive and fine-grained exploration of the predicted protein structure space while maintaining interactive performance at scale.

### Description of the Web server

AlphaFind v2 is a web application allowing interaction via a browser with a backend primarily executing alignment computations, and a vector database, which performs approximate search.

#### Data preparation

##### Embeddings creation

AlphaFold DB (v4) was downloaded and each protein structure compressed into 1536-dimensional embeddings using the ESM3 generative model combined with a transformer neural network from (10). In addition to whole-chain structure embeddings, embeddings were also computed for all structures after removal of unstable regions (pLDDT *<* 70 / 80 / 90). For TED domains, pre-computed Foldclass 128-dimensional embeddings (6) were downloaded from (5).

##### Metadata extraction

All the relevant metadata – such as organism, taxonomy ID, gene name and protein name – were extracted from every protein structure file. For TED domains, the metadata provided by (5) were downloaded, which included the residue boundaries identifying the domain within a protein.

##### Vector database ingestion

The embeddings and metadata were then stored within the OpenSearch vector database, each search mode within its own index using the HNSW index with 16x compression and running in the “on disk” mode.

### Workflow

The AlphaFind v2 includes six alternative search modes: full-chain, pLDDT-filtered chains (at thresholds 70, 80, 90), and domains (*TED, TED Multidomain*). Its search workflow consists of three phases.

In Phase 1, the user query is validated and converted into a fixed-length embedding using the embedding pipeline described in the previous section. In Phase 2, this embedding is used by the vector database to find the most similar embeddings via approximate k-nearest neighbor search (*k* = 100) with cosine distance. This step outputs a ranked list of candidates, which is returned to the user immediately. Phase 3 runs in the background to refine structural similarity ranking without blocking the user interface. It involves the computation of pairwise alignments between the query structure and the candidates using US-align (15), producing TM-score, structural alignment and RMSD. Consequently, while the ranking order may shift, the underlying set of candidates remains unchanged between Phase 2 and Phase 3. The last phase also links any known experimental structure with the retrieved results. The user can optionally choose to load more structures than the default 100, in which case the candidate list is extended and Phase 3 repeats. The process is visualized in Figure 1. For details about the TED domains workflow, see Supplementary Text S1.

**Figure 1.**
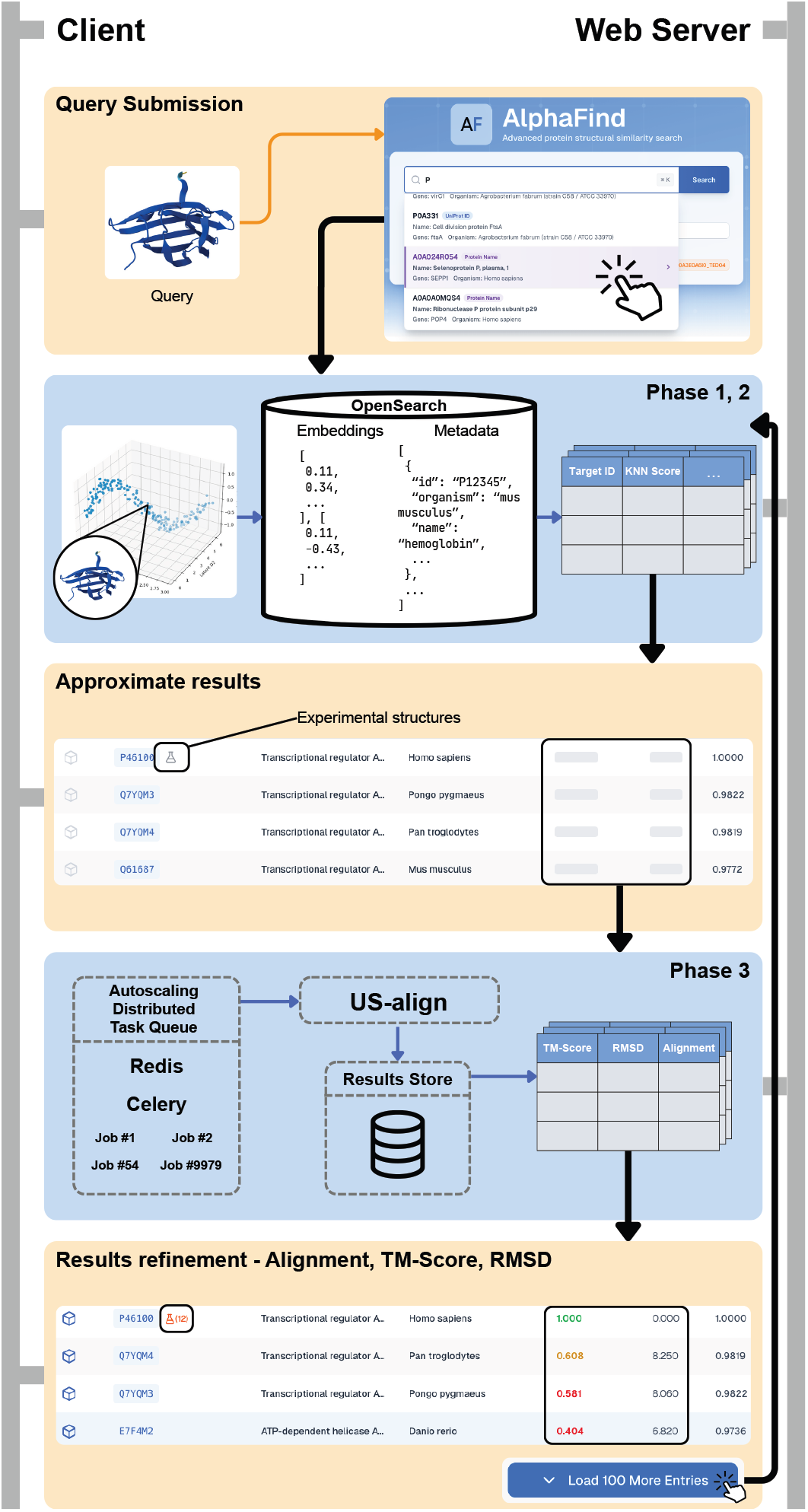
Search workflow of AlphaFind v2.

### Implementation

The backend is implemented in Python as a modular service-oriented stack: a Flask-based REST API handles request validation and dispatch, while long-running computations (TM-score refinement, experimental structure resolution) are executed asynchronously using Celery workers coordinated via Redis to minimize latency. Vector embeddings and metadata are stored and queried in OpenSearch, with separate indexes per search mode to ensure predictable retrieval behavior. PostgreSQL is used for persistent state management of queries, task lifecycles, and server metadata, and also acts as a caching layer for frequently accessed results. The stack is deployed on Kubernetes, enabling horizontal scaling of computational workers to dynamically accommodate refinement workloads and API requests.

### Interface / Interaction

The search results are organized into seven tabs. The first six are dedicated to the AlphaFind v2 search modes: full-chain, pLDDT thresholds of 70, 80 and 90, TED domains, and TED Multidomain. The last tab includes the legacy AlphaFind v1 search (9). This mode is most akin to AlphaFind v2 full-chain mode, however, it uses a different embedding (8) and approximate search method (1), and thereby produces different sets of results.

In each tab, the results are shown in a table, each row corresponding to one chain, domain, or a multidomain aggregation found to be a match. The table contains columns including UniProt ID, Protein Name, Organism Name, and similarity measures: TM-score, RMSD, and kNN Score, the latter referring to the cosine similarity of the protein embeddings. The results are sorted by TM-Score; however, while Stage 3 (refinement of the ranking) is still running, the kNN Score is used as a surrogate. Each row also contains a button to trigger 3D alignment, which displays both query and target structures in an embedded Mol* viewer (11) inheriting its full functionality. The alignment, computed by US-align, is only taking into account residues relevant for the search mode (e.g., only computed on the specific domain residues in case of TED). The visualization distinctly shows which residues from the chain are considered and which are not.

In the TED Multidomain mode, the user gets additional fine control on the alignment in the form of sliders for each of the domain pairs, to specify its contribution to the overall alignment. As the user adjusts the sliders, the alignment of the structures in the 3D view changes interactively to prioritize a given domain-target pair. This functionality allows users to gradually shift focus from a detailed inspection of a single domain pair, through the interaction of several pairs, to the global alignment of the whole structures.

### Limitations

AlphaFind v2 uses the AlphaFold DB version 4, as the most recent version 6 is not available for bulk download at the time of publication, with the exception of some selected proteomes.

The search uses a vector database with approximate search to retrieve the relevant embeddings. This search strategy was chosen for its speed, but it does not guarantee to find the objectively best answer. However, the application’s design allows for extending the results set, which improves the prior search answer.

## Results and discussion

### Performance

To assess AlphaFind’s performance, we consider two metrics: (i) the time it takes to retrieve the results and (ii) the average TM-Score (16) of the top-n retrieved results. We used a query set composed of 2050 multidomain proteins from (10) to test the *full-chain* search mode. From the same protein set we extracted 4420 TED domains and used these in the *domains* search mode.

Table 1 shows that AlphaFind v2 finds the approximate answer an order of magnitude faster than the other methods. TM-Score refinement is finished faster than AlphaFind v1 search and comparable to the FoldSeek Server search time. In domains search mode, AlphaFind v2 is significantly faster than Merizo-search. The results also show that the speedup comes with statistically superior TM-Score at the *p <* 0.05 level across all comparisons. FoldSeek does not return TM-Scores, these were computed separately as part of our evaluation and not counted in the measured time. FoldSeek also operates on the AlphaFold database clustered at 50% sequence identity (2) and its greater structural diversity can skew TM-score results in favor of AlphaFind, which uses the raw AlphaFold database. Merizo-search uses the same TED database as AlphaFind v2.

**Table 1.** Runtime and retrieval performance comparison of AlphaFind with FoldSeek Server and Merizo-search. *All* corresponds to full retrieval (≈ 800 results for chains, ≈ 100 for domains). For AlphaFind v2 we report two sets of times – approximate, *“a”*, denoting the time taken to get the approximate results on the embeddings and *“tm”*, marking the time it takes to get the refined results, including alignment, TM-score and RMSD on the raw structures.

| Method | Time<br>*mean* ± *std*<br>(s) | Avg.<br>TM-Score<br>(top 10) | Avg.<br>TM-Score<br>(top 100) | Avg.<br>TM-Score<br>(all) |
| --- | --- | --- | --- | --- |
| *Chains* |  |  |  |  |
| AlphaFind v2 | 2.40 ± 1.89 (a)<br>45.88 ± 54.27 (tm) | 0.733 | 0.654 | 0.588 |
| AlphaFind v1 | 93.38 ± 36.65 | 0.670 | 0.546 | 0.364 |
| FoldSeek Server | 42.03 ± 37.47 | 0.596 | 0.532 | 0.440 |
| *Domains* |  |  |  |  |
| AlphaFind v2 | 0.49 ± 0.41 (a)<br>31.28 ± 31.11 (tm) | 0.947 | 0.894 | — |
| Merizo-search | 145.97 ± 38.90 | 0.865 | 0.824 | — |

### Case studies

#### PIN3 auxin carrier protein, “pLDDT ≥ 90” search mode

PIN auxin carrier proteins are essential transmembrane transporters that regulate plant growth by moving auxin from the cytosol to the extracellular space, categorized as “long-loop” or “short-loop” based on the size of their intrinsically disordered cytosolic loop. Due to this high level of disorder, experimentally determined structures such as PIN3 (12) contain large unstructured segments that complicate traditional structural homology searches. When searched in the *full-chain* search mode, no PIN3 homologue has a TM-score above 0.7 and many PINs from soybean are omitted. As shown in Figure 2, AlphaFind v2 is able to focus searches exclusively on stable, high-confidence regions (pLDDT *>* 90), with a hit of TM-score 0.947 and find multiple soybean PIN proteins. This enables the efficient identification and analysis of long-loop PIN proteins across diverse plant species by filtering out structural noise.

**Figure 2.**
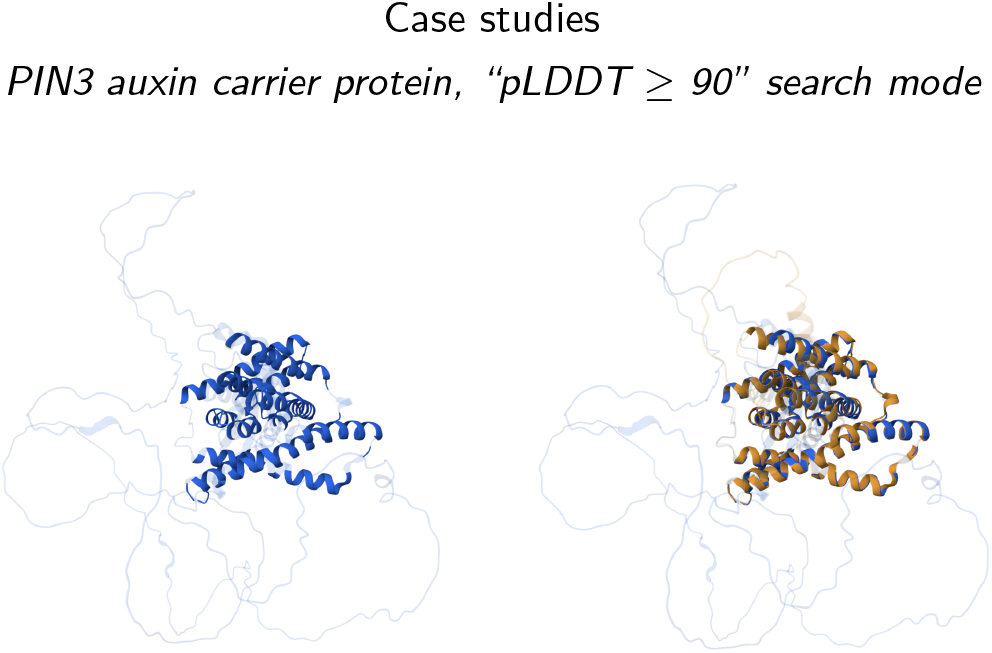
Left: PIN3 auxin carrier protein from *Arabidopsis thaliana* (UniProt ID: Q9S7Z8). Right: Q9S7Z8 (blue) aligned with PIN3 auxin carrier homolog from soybean (UniProt ID: A0A0R0ILJ6, orange). Brightly colored residues have pLDDT *>* 90, and are considered in the alignment computation.

#### NCAM1, “TED Multidomain” search mode

Neural Cell Adhesion Molecule 1 (NCAM1) is a cell surface glycoprotein involved in neuronal adhesion, synapse formation, and nervous system development. It belongs to the immunoglobulin superfamily and is conserved across vertebrates. The NCAM1 ectodomain has a modular architecture composed of five immunoglobulin domains followed by two fibronectin type III domains.

These domains occur in many other proteins. However, their specific order and combination are characteristic for NCAM1. In this use case, the *multidomain* search mode captures this domain arrangement rather than individual domains (Figure 3, top part).

**Figure 3.**
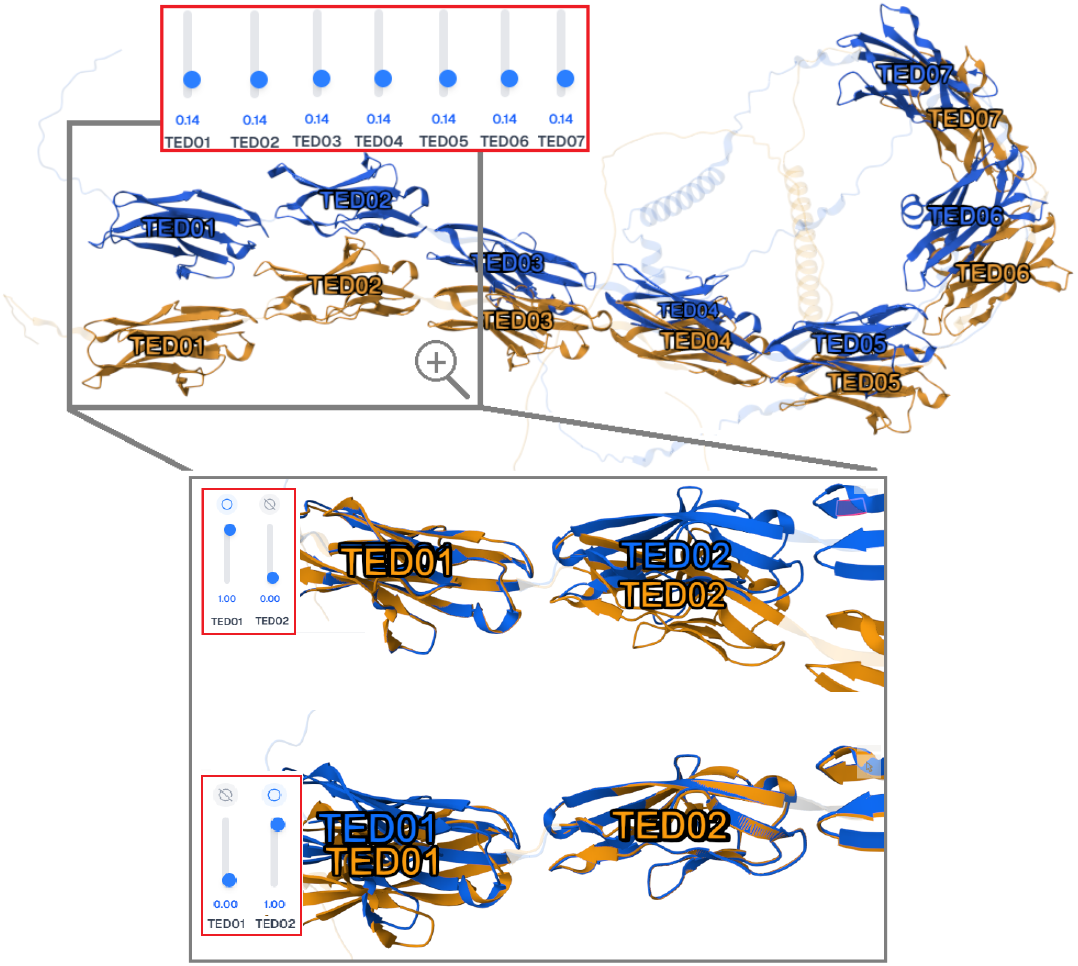
Structural alignment in the TED Multidomain search mode for NCAM1. Top: *Homo Sapiens* (UniProt ID: P13591, blue) aligned with its *Felis Catus* homolog (UniProt ID: Q5G7G8, orange). Brightly colored residues represent the TED domains used for alignment computation. The toggle weights are set equally to reflect uniform domain contribution to the overall alignment. Bottom: Weighting the alignment of individual TED domains – TED01 and TED02 – in the TED Multidomain analysis panel.

This enables the identification of other adhesion proteins with a similar multidomain architecture and highlights structural and evolutionary relationships within the Ig superfamily.

This case study also demonstrates the usefulness of complex alignment of 3D structures. With most matching targets, if one of the matching domain pairs is perfectly aligned, most of the other pairs become misaligned due to their different positions in the distinct AlphaFold predictions. On the other hand, if the alignment is averaged, the individual pairs are not sufficiently aligned to assess their detailed visual similarity. Therefore, complex manipulation of the alignment weights is necessary to tune the desired view (see Figure 3, bottom part).

## Conclusion

In this article, we present AlphaFind v2, a web application for fast structure-based search of similar proteins in the AlphaFold Protein Structure Database. AlphaFind v2 extends the first version of the application (9) by supporting multiple complementary search modes, including full-chain, confidence-filtered, domain, and multidomain structural comparisons. The application combines embedding-based pre-filtering with alignment-based refinement and presents the results using quantitative similarity metrics, rich metadata, and interactive 3D structural superpositions. AlphaFind v2 is easy to use, platform-independent, and together with its detailed documentation, freely available at https://alphafind.ics.muni.cz/.

## Supporting information

Supplementary Text S1: Workflow - TED domains

## Conflicts of interest

The authors declare that they have no competing interests.

AlphaFind v2: Similarity Search in AlphaFold DB and TED Domains across Structural Contexts, YEAR, Volume XX, Issue x

## Data availability

AlphaFind v2 is free and open to all users at https://alphafind.ics.muni.cz/ and there is no login requirement.

The data (pre-computed protein embeddings and metadata) is available at https://doi.org/10.58074/mv1w-y227.

## Author contributions statement

Terézia Slanináková (Conceptualization, Methodology, Software, Data processing, Writing—original draft, Evaluation), Adrián Rošinec (Software, Writing—original draft), Jakub Čillík (Software, Visualization), Aleš Křenek (Conceptualization, Methodology, Writing—original draft), Katarina Gresova (Conceptualization, Evaluation, Testing), Jana Porubská (Writing—original draft, Testing), Eva Maršálková (Data processing), Jaroslav Olha (Conceptualization, Writing—review & editing), David Procházka (Testing, Writing—review & editing), Lukáš Hejtmánek (Software —Infrastructure), Vlastislav Dohnal (Funding acquisition, Writing —review & editing), Karel Berka (Testing, Writing—review & editing), Radka Svobodová (Funding acquisition, Supervision, Writing—original draft), Matej Antol (Conceptualization, Funding acquisition, Supervision, Writing—original draft).

## Acknowledgments

The authors thank the anonymous reviewers for their valuable suggestions. The authors also thank Tomáš Pavlík, Jozef Sabo, Viktória Spišáková, Michal Mikuš, Tomáš Svoboda and David Sehnal for their help related to this publication.

