## Supplementary Text S1: Workflow - TED domains for "AlphaFind v2: Similarity Search in AlphaFold DB and TED Domains across Structural Contexts"

In the TED domains search mode, the query and the candidates are protein domains. In the TED Multidomain mode, the results of search for all the query domains are merged. For each target, a combined, altered TM-score is computed:

$$TM_{\text{target}} = \frac{1}{N_{\text{target}}} \sum_{i=1}^{N_{\text{common}}} \frac{1}{1 + d_i^2} \tag{1}$$

where $N_{\text{target}}$ is the number of domains in the target protein ($TM_{\text{query}}$ can be defined symmetrically), $N_{\text{common}}$ is the number of domains which have their matching counterparts in both query and target, and $d_i$ is RMSD between the $i$-th pair of matching query-target domains after their independent alignment. This TM-score behaves as one would expect – it grows higher with better match of individual domains as well as with more domains considered.
